# Computer Simulation of the interaction between SARS-CoV-2 Spike Protein and the Surface of Coinage Metals

**DOI:** 10.1101/2022.07.28.501856

**Authors:** Mehdi Sahihi, Jordi Faraudo

## Abstract

A prominent feature of the SARS-CoV-2 virus is the presence of a large glycoprotein spike protruding from the virus envelope. The spike determines the interaction of the virus with the environment and the host. Here, we used an all-atom molecular dynamics simulation method to investigate the interaction of up and down conformations of the S1 subunit of the SARS-CoV-2 spike with the (100) surface of Au, Ag and Cu. Our results revealed that the spike protein is adsorbed onto the surface of these metals, being Cu the metal with the highest interaction with the spike. In our simulations, we considered the spike protein in both its up conformation S^up^ (one receptor binding domain exposed) and down conformation S^down^ (no exposed receptor binding domain). We found that the affinity of the metals for the up conformation was higher than their affinity for the down conformation. The structural changes in the Spike in the up conformation were also larger than the changes in the down conformation. Comparing the present results for metals with those obtained in our previous MD simulations of S^up^ with other materials (cellulose, graphite, and human skin models), we see that Au induces the highest structural change in S^up^, larger than those obtained in our previous studies.

**GRAPHICAL ABSTRACT:** 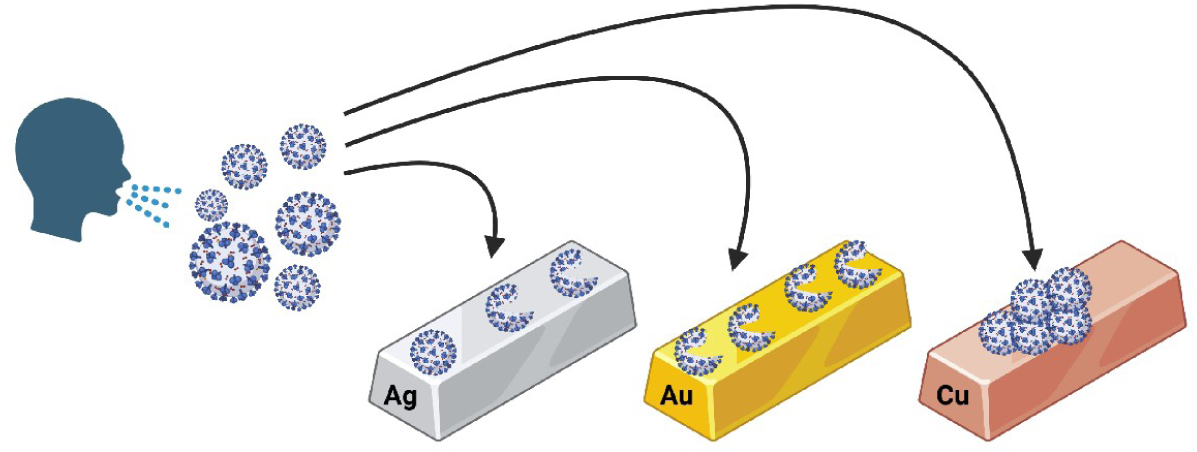

## 1. INTRODUCTION

Many types of viruses, especially respiratory viruses such as SARS-CoV-2, are spread by virus-containing aerosols and droplets which can contaminate surfaces. The potential presence of infective viruses on the surfaces of materials is the main reason for recommendations of the health authorities on frequently disinfecting surfaces and washing hands. An interesting alternative to prevent surface-mediated indirect transmission of viruses is to design virucidal surfaces/coatings that can be used in highly touched surfaces [1]. In practice, this may add initial costs, and also there are significant difficulties in assessing the virucidal activity of materials [2]. But development of antiviral materials or coatings could play a crucial role in inhibiting the spreading of present day viruses and possible future pandemics. Development of the virucidal materials can also supplement the available cleaning and disinfection strategies and create a comprehensive defense against indirect transmission of viral diseases.

The antiviral activities of different types of materials, i.e., metals (silver, copper and gold), metal oxides (TiO_2_ and Fe_2_O_3_), clays, carbon-based materials (fullerenes and carbon nanotubes) and their mixtures has been previously studied in the literature [3–9]. There are many mechanisms contributing to the potential virucidal activity of a material. Some of them are ion release (as in the case of clays or in materials containing metals [9]), chemical reactions that can generate reactive oxygen species that damage the virus [1] and disruption of the virus envelope due to interactions between virus components and surfaces [5, 7]. In general, the stability of viruses can be changed substantially (either increased or decreased) due to their adhesion to surfaces, and this change depends on both the material and environment conditions [10, 11].

An illustrative example of the dramatic effect of the material is provided by recent studies on SARS-CoV-2 [12]. The virus remains infectious up to 72 h after application on plastic and stainless-steel surfaces, while it is inactivated after 4 h on a copper (Cu) surface. Furthermore, inactivated (non-infective) SARS-CoV-2 viruses showed even shorter stability on a Cu surface [13]. This behavior of Cu for the SARS-CoV-2 case is consistent with the accumulated historical evidence of Cu as a biocidal agent [14–17]. Furthermore, the virucidal effect of Cu against Phi6, influenza virus, norovirus, monkeypox, vaccinia virus, HIV, SARS-CoV, and SARS-CoV-2 has been studied, before [18–25]. The US Environmental Protection Agency (EPA) has also approved the antimicrobial effect of more than 500 Cu alloys [14, 17, 26–28]. Therefore, Cu can be considered a nontoxic, economical and resource-efficient substance against the spread of pathogens [26, 27].

It is reasonable to think that this behavior of Cu may be shared with other metals. In fact, the biocidal effect of the coinage metals Ag, Au and Cu has been known for long, being, first described systematically by Karl Nägeli in 1893 [28]. In the case of Ag, it is known that an Ag solid surface can bind to viral surface proteins and/or denature enzymes, inhibiting the action of different types of viruses [29, 31]. Also, Ag is being increasingly used as an effective antimicrobial agent in wound care harvests, medical strategies, textiles and water disinfecting filters [31–40]. Moreover, silver nanoparticles (AgNPs) inhibit various types of viruses including herpes simplex virus, human parainfluenza virus type 3, hepatitis B virus, etc [41–49]. Furthermore, Han et al. have proven antiviral activity of Ag/Al_2_O_3_ and Cu/Al_2_O_3_ surfaces against SARS-CoV-1 coronavirus and baculovirus [50].

As it was mentioned before, Au is also another metal with biocidal activity. It has been suggested that AuNPs attach onto the virus’s glycoprotein to hinder virus performance [51]. Lysenko et al. showed that silicon dioxide-coated AuNPs have almost a 100% inhibitory effect against adenovirus [52]. Also, a new Au nanorod has been developed against the MERS virus [53]. Modified and non-modified AuNPs can inhibit HSV-1, HSV-2, HIV and influenza-A viruses [54–58]. Furthermore, Knez et al., have investigated the adsorption of Tobacco Mosaic Virus onto Au surface using AFM method [59]. All this experimental evidence motivates the relevance of the study of the interaction of viruses with metal surfaces [60–62].

Generally, viruses are adsorbed onto the mineral and charged surfaces via vdW and electrostatic interactions, respectively [63–66]. As viruses tend to be more hydrophobic than proteins, it is suspected that vdW and hydrophobic interactions have dominant roles at the interface of the virus-metal surface [67, 68]. In any case, knowledge of the virus-surface interaction is still rather limited. The lack of a fundamental knowledge of the physico-chemical aspects of the virus interaction with metals is in contrast to the available atomistic detailed information about the different types of viruses. SARS-CoV-2 virus has the typical structure of a coronavirus, and its glycoprotein spikes are responsible for the virus interaction with host cell receptors and the environment. Therefore, in the present study we investigate the interaction of fully glycosylated spike protein (S^pro^) of SARS-CoV-2 with surfaces of different metals (Ag, Au, and Cu). All atom molecular dynamics (MD) simulation method is used to clarify the mechanism, molecular and atomic details of these adsorptions and also to complement our previous works on the interaction between SARS-CoV-2 and different types of hard and soft materials [69, 70]. Furthermore, because of the presence of two different conformations (up and down states) for S^pro^ [71, 72] and their equal proportion on the surface of virion [73], the interactions of both of the conformations with the metal surfaces are investigated and compared. Most of the published MD simulation studies have only investigated the interaction of the up-conformation of S^pro^ (S^up^) with the surface of the materials, and very rare studies have considered the interaction of down-conformation (S^down^), as well [74].

## 2. MATERIALS AND METHODS

### 2.1. System preparation

S^pro^ consists of three identical polypeptide chains and is divided into the S1 (residues 1-1146 per chain) and S2 (residues 1146-1273 per chain) subunits. The S1 subunit is the head of the protein that is exposed to the exterior and is responsible for contact with metal surfaces. However, the S2 subunit links the protein to the virion. Fully glycosylated structures of the S1 subunit of S^pro^ were taken from CHARMM-GUI archive [75] (PDB IDs: 6VSB and 6VXX for S^up^ and S^down^, respectively, **Figure S1**). The difference between the S^up^ and S^down^ is based on orientation of their receptor binding domain (RBD). The downloaded structures are based on cryo-EM resolved crystal structures, reported by Walls et al. [72], and have the predicted missing residues and the linked glycans reported by Woo et al. [76]. The binding glycans may affect the interaction of S^pro^ with the metal surfaces. S^pro^ consists of 165 glycans per subunit. The obtained structures contain 72990 atoms and their total charge (at pH=7) is −15e. S^pro^ structures were solvated using the “gmx editconf” and “gmx solvate” modules of gromacs [77–80] in a cubic box, and then all the water molecules beyond 3 Å of a solvation shell were removed. The number of water molecules added to solvate glycosylated S^pro^ were 83809 and 67041 for S^up^ and S^down^, respectively. We also neutralized the systems by adding K and Cl ions at a concentration of 150 mM.

The Avogadro software [81] was used to prepare the structure of Ag (100), Au (100) and Cu (100) slabs with dimensions of 30.00 × 30.00 × 1.20 nm^3^. Ag and Au slabs consist of 69312 atoms in six layers and Cu slab consists of 98784 atoms in seven layers.

To study the wetting behavior of the metals, a water droplet with a diameter of about 0.75 nm (6845 water molecules) was generated using VMD software [82]. The solvated and neutralized structures of S^pro^ and also the water droplets (for wetting calculations) were placed on top of the metal slabs and their distance was set to approximately 5.0 Å (**Figures 1** and **S1**). As shown in these figures, the initial orientation of the spike was such that the long axis of the spike was perpendicular to the surface, mimicking the expected orientation for the spike of a virus approaching to the surface. The total number of atoms for each system is mentioned in **Table 1**.

**Figure 1.**
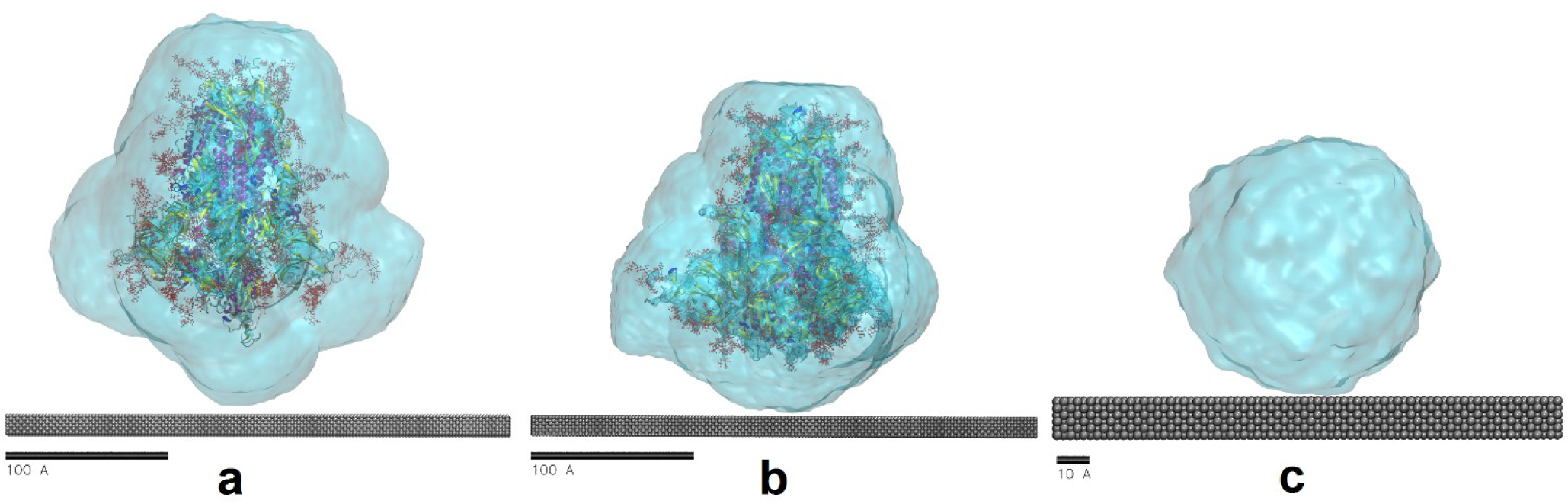
The solvated and neutralized structures of S^pro^ (a and b for S^up^ and S^down^, respectively) and also the water droplet (c; for wetting calculations) on the top of the Ag slab (distance was set to approximately 5.0 Å). For the Au and Cu slabs we have similar starting structures.

**Table 1.** The total number of atoms for S^pro^-metal and water-metal systems.

| System | Ag-S<sup>up</sup> | Au-S<sup>up</sup> | Cu-S<sup>up</sup> | Ag-S<sup>down</sup> | Au-S<sup>down</sup> | Cu-S<sup>down</sup> | Ag-Water | Au-Water | Cu-Water |
| --- | --- | --- | --- | --- | --- | --- | --- | --- | --- |
| No. atoms | 386026 | 386026 | 415498 | 335532 | 335532 | 365004 | 37863 | 37863 | 45231 |

### 2.2. MD simulation

All the MD simulations were done using the simulation package GROMACS version 2020.5 [77–80] for S^pro^-metal and water-metal complexes. The CHARMM36 force field was employed in all the simulations. This force field considers parametrization of carbohydrate derivatives, polysaccharides, and carbohydrate–protein interactions [83]. The TIP3P water model included in CHARMM36 is also used in our simulations. Most of the force fields, including CHARMM36, do not have parameters for all of the elemental metals we are studying and we must choose the level of accuracy that can be considered for Ag, Au and Cu. Therefore, like some of the previously published MD simulation studies [84–86] we prefer to model the metal surfaces at the classical level, using the Lennard-Jones (LJ) potential (**Table 2**). Previous studies showed that interfacial water structure has not been influenced by polarization of the force field [87]. On the other hand, not only the polarization portion is less than 10% of the total energy [88] but also the charges interact with their images, and decrease the polarizability influence [89, 90]. Considering the limitations of our model, we characterize the wetting behavior of the metal surfaces to verify the accuracy of the used model. Concerning the size of the metal slab to be simulated, we consider six and seven atomic layers for Ag/Au and Cu, respectively, which should be enough due to the short range of the LJ potential.

**Table 2.** Nonbonded force field parameters used for metals.

| Metal | $\sigma/\text{nm}$ | $\epsilon/\text{kJ.mol}^{-1}$ | $\epsilon/\sigma$ |
| --- | --- | --- | --- |
| Ag | 0.26326057121 | 19.07904 | 72.47207553 |
| Au | 0.26290421172 | 22.13336 | 84.18792478 |
| Cu | 0.23305910466 | 19.74848 | 84.73593011 |

The systems consisting of a solvated S^pro^ and metal slabs of Ag, Au or Cu were placed in the center of a cubic box, and the minimum distance between the system and the box boundaries was set to 1.0 nm. All layers of the heavy metal slabs were geometrically frozen during the simulations to approximate a realistic metal slab configuration. Integration of the equations of motion was done at a time step of 2 fs with full periodic boundary conditions (PBC) applied along the three Cartesian directions. The systems were energy minimized using the conjugate gradient (CG) method, with 1×10^−6^ (kJ.mol^−1^ and kJ.mol^−1^.nm^−1^ for energy difference and RMS force, respectively) convergence criteria. Then, we performed 100 ns of NVT production runs (200 ns for adsorption of S^pro^ in its up state onto the Cu surface, this system was equilibrated later than the other systems) at 300 K using a Berendsen thermostat [91] with a damping constant of 0.1 ps. Also, 2 ns of NVT productions were done for wetting calculations. The Berendsen thermostat has been widely employed in previous simulation works of proteins, showing good agreement with experiments [92–95]. During the simulations, a 1.0 nm cutoff for LJ and Coulomb interactions was applied and the particle mesh Ewald method [96, 97] was used for long-range electrostatics.

The LINCS method [98] was also used as a constraint algorithm. All the images were generated using VMD software [82].

## 3. RESULTS AND DISCUSSION

### 3.1. Wetting behavior of the metal surfaces

To verify the accuracy of the used model and force field parameters for system components, we characterized the wetting behavior of the metal surfaces by placing a water droplet on top of them. As shown in **Figure S2,** the average root mean square deviation (RMSD) values of the metal-water systems are about 4.18±1.03, 4.30±1.06, and 5.23±1.46 nm for Ag, Au and Cu surfaces, respectively. In fact, the RMSD of the system reached equilibrium and fluctuated around its mean values after about 1 ns indicating that the system was well-behaved thereafter and could be analyzed in its equilibrium state to calculate the equilibrium contact angle. **Figure 2** shows the final configuration of water droplets on the surface of the studied metals. The analysis of the results showed almost full wetting (or a small contact angle) for all the three metals. This behavior has also previously been reported using MD simulation and experimental methods [99–101]. Hence, it could be concluded that the use of the TIP3P water model in combination with the nonbonded force field parameters for metals employed here, is in agreement with experiments and they can be used in our simulation of the adsorption of hydrated S^pro^ onto the metal surfaces. On the other hand, the configuration of water molecules on the metal surfaces shows that there are two distinct layers of water molecules on the surfaces: the interfacial (first) layer and the bulk water. This observation has also been reported and investigated in detail by other groups [102, 85]. This could be considered as another evidence to prove the appropriateness of the used parameters for our MD simulation studies.

**Figure 2.**
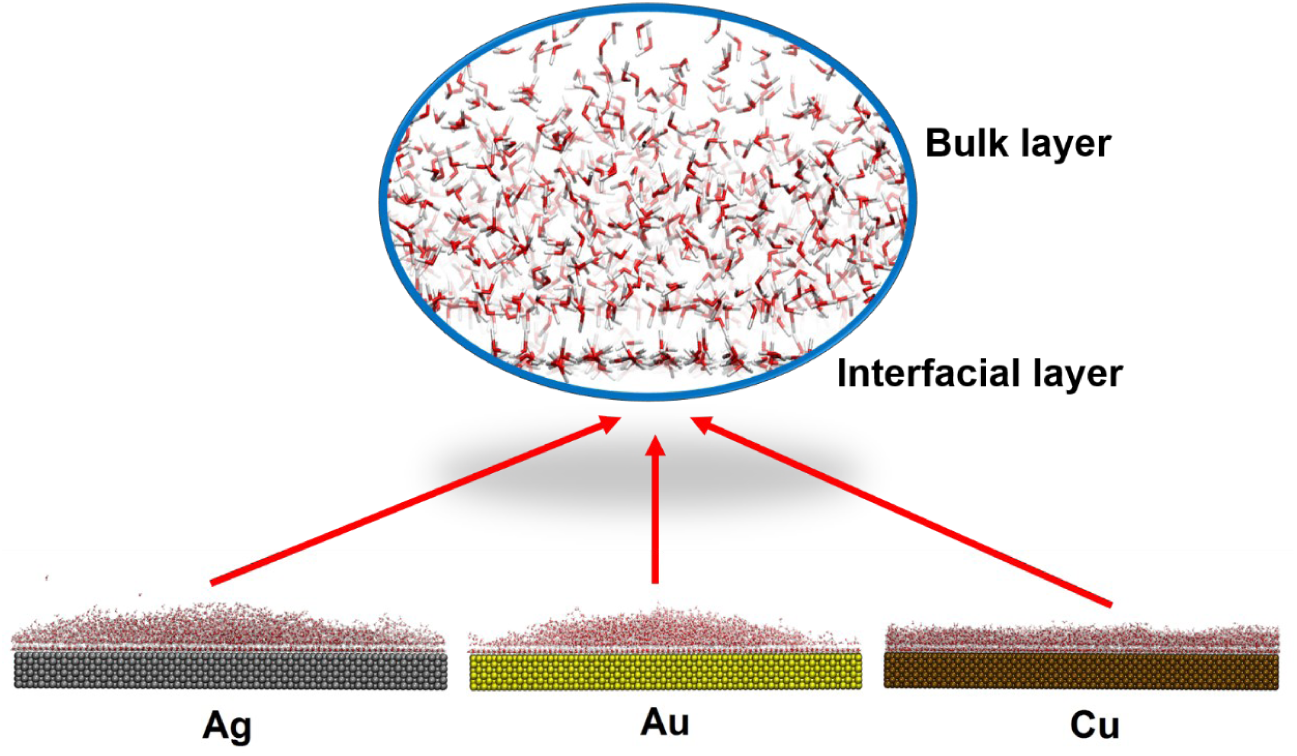
Final configurations of water droplet on the surface of Ag, Au and Cu.

### 3.2. Adsorption of S^pro^ onto the surface of metals

#### 3.2.1 S^up^ conformation

The trajectories show that S^up^ is adsorbed onto the surfaces of investigated metals with the almost similar mechanism: the protein adjusted its spatial conformation in a few time steps, then started to interact with the surfaces rapidly, and finally achieved the comparatively equilibrium state with readjusted conformation. However, the final conformations of the protein on the metal surfaces are completely different in each case. **Figure 3** shows snapshots of the final configuration of S^up^ on the metal surfaces. As shown in **Figure 3** (top image), S^up^ started to contact the Ag surface at about 10 ns by its glycan groups, and finally were adsorbed on the Ag surface with more contacts at 100 ns. For Au, the contact of S^up^ started at 2 ns by the glycans and its final structure shows more deformation in comparison to Ag surface (**Figure 3**, middle image). In the case of Cu, the contact of S^up^ started even earlier than Au but shows less tilt angle in comparison to two other metals (**Figure 3**, bottom image). The solvation water is not shown in the figure to simplify the visualization (**Figure S3** includes solvation water).

**Figure 3.**
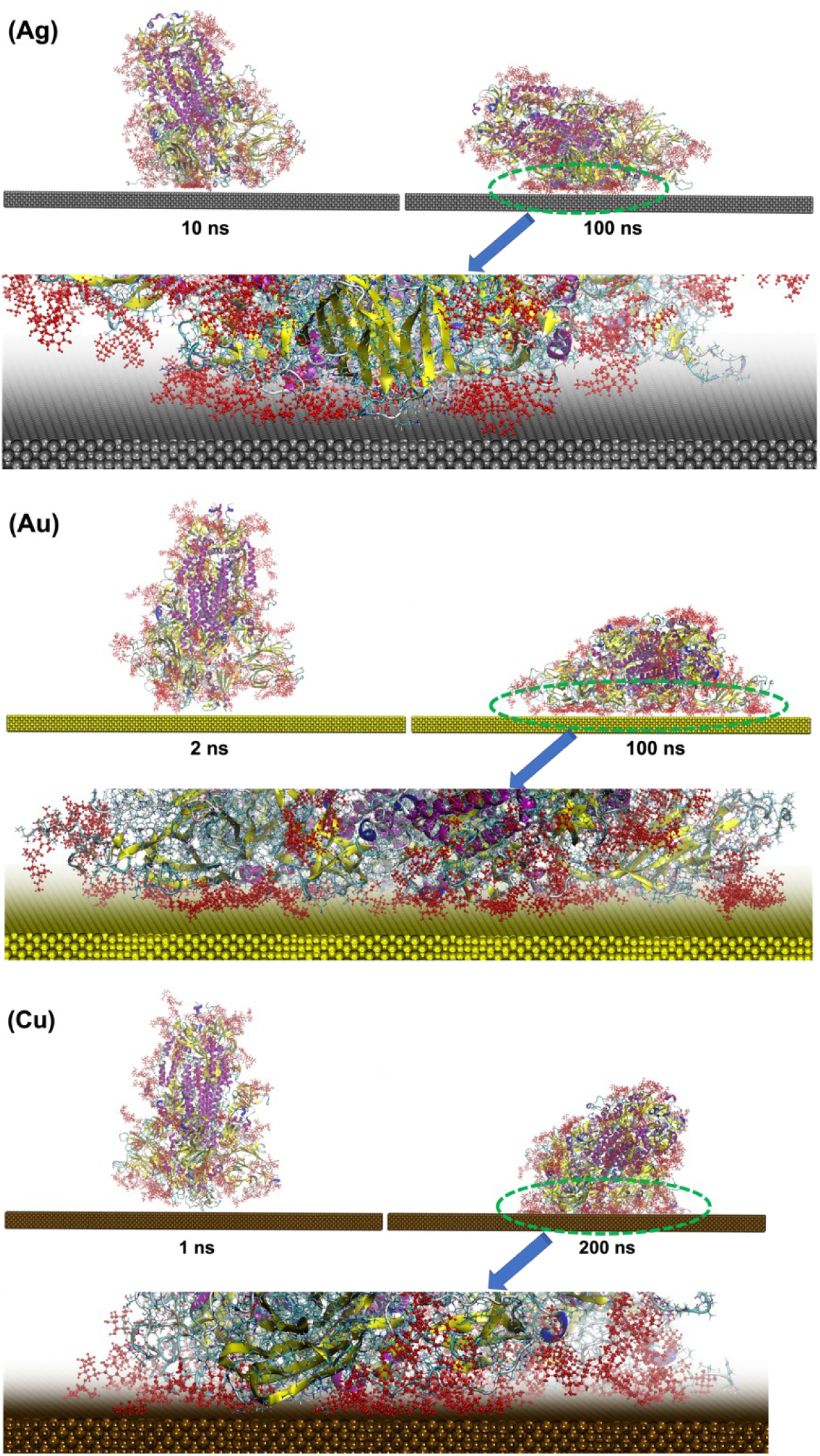
Representative snapshots of S^up^ adsorbed onto Ag, Au, and Cu surfaces. The number and type of amino acid residues (in the final time frame of the trajectories) in contact with metals are shown in Figure 4. The protein is shown in CPK representation (with its structure emphasized in cartoon representation). S^pro^ glycans are shown in red. Water is not shown for simplicity.

A quantitative analysis of the interaction between S^up^ and the metal surfaces along the simulated trajectories was done by computing the following magnitudes: the contact area between S^up^ and the metal surfaces, the number of protein residues in contact with the surfaces and the LJ interaction energy between S^up^ and the metal surfaces. The contact area (**Figure 4a**) is calculated as [103]:

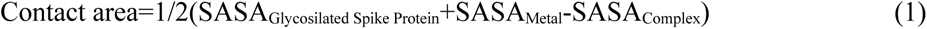

where SASA is the solvent-accessible surface area. The contact area obtained for Au is about three times larger than the ones obtained for Cu or Ag, as shown in **Figure 4** (see also the snapshots of **Figure 3**).

**Figure 4.**
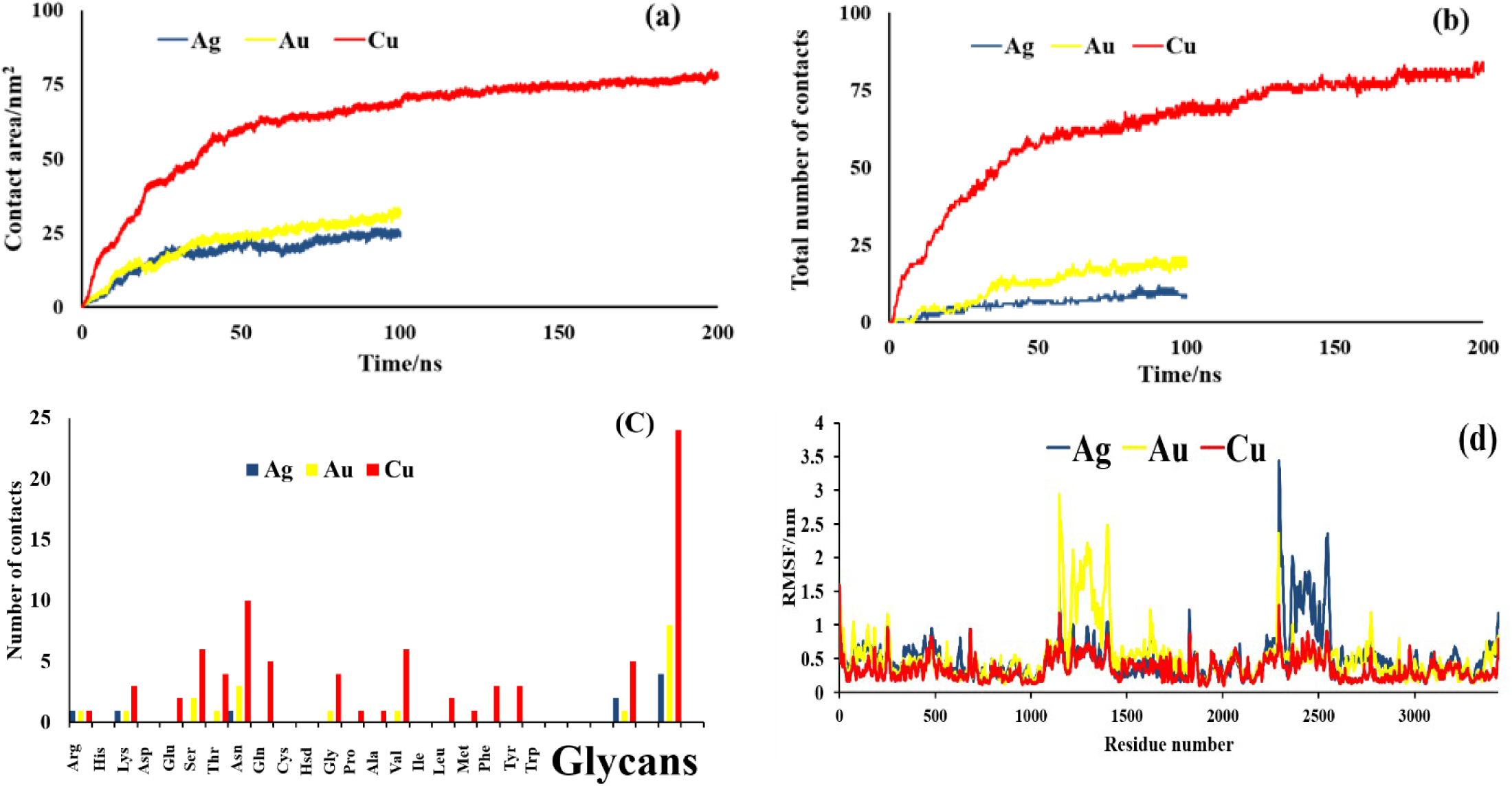
(a) Contact area between S^up^ and metal surfaces as a function of time, (b) Total number of S^up^ residues in contact with the metal surfaces as a function of time, (c) Average number of S^up^ amino acids (3-letter code) in contact with the metal surfaces at equilibrium, and (d) RMSF of S^up^ amino acid residues.

We have also computed the total number of amino acid residues in contact with the metal surfaces by considering that the surfaces are in contact with one amino acid if they have at least one pair of atoms separated by a distance not larger than 3.5 Å as in our previous works [70].

The total number of residues in contact with the Cu surface (**Figure 4b**) is larger than Ag or Au. Also, its evolution with time shows the same behavior as the contact area (**Figure 4a**). Indeed, these observations suggest that the affinity of S^up^ to the metal surfaces follows the sequence Cu >> Au > Ag. As it was mentioned in the introduction section, vdW and hydrophobic interactions have dominant roles at the interface of virus-metals [67, 68]. In this regard, **Figure 4c** represents that amino acid residues with polar uncharged side chains (Ser, Thr, Asn and Gln), amino acid residue with hydrophobic side chain (Val), and glycan groups, which responsible for vdW and hydrophobic interactions, stabilize the metal-S^up^ complexes (especially in the case of Cu).

The residue-based root-mean-square fluctuation (RMSF) is calculated based on average positions of amino acids to evaluate their local dynamical variation and identify the regions of the protein that have high structural changes and fluctuations during the simulation. As shown in **Figure 4d**, the first and last amino acid residues (the N- and C-terminals) of each S^up^ monomer have a high RMSF due to their inherent high flexibility. Furthermore, the RMSF values for the residues of S^up^ in contact with Cu surface are lower than the RMSF values for protein in contact with Ag and Au surfaces. It means that the combination of S^up^ with Cu stabilizes the protein and causes less flexibility of its amino acid residues.

**Figure 5** shows the evolution of the interaction energy between S^up^ and the metal surfaces as a function of time. The changes in LJ interaction energy are in agreement with the above-mentioned behaviors of contact area and total number of contacts. Indeed, when S^up^ is adsorbed onto the Ag and Au surfaces (0-30 ns of the simulation time), the interaction energy decreases and then remains almost constant until the end of the simulation. The obtained final values were about −2313.51±240.38 and −3210.98±355.27 kJ.mol^-1^ for Ag and Au, respectively, which indicate a stronger interaction of S^up^ with Au as compared with Ag, consistent with the higher contact surface and higher number of amino acids in contact.

**Figure 5.**
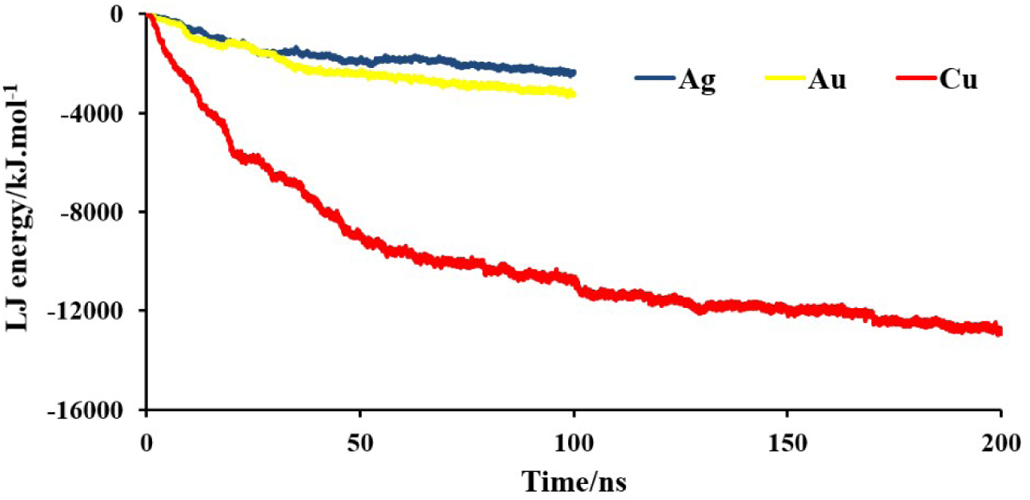
LJ interaction energy between S^up^ and the metal surfaces as a function of time.

For Cu, the decrease in interaction energy and increase in contact area evolved rapidly over time until about 50 ns and remained stable at about −12780.50±1532.92 kJ.mol^-1^ and 77.95±5.49 nm^2^ for LJ interaction energy and contact area, respectively.

These results can be interpreted by noting that the affinity of metals for S^up^ is directly correlated with the metal LJ parameters (see **Table 2**). For example, Cu that shows the highest interaction energy with S^up^ has the highest ε/σ ratio and the lowest ε value among the studied metals. Indeed, the affinity of S^up^ for the metals is in agreement with their ε/σ ratio and follows the sequence Cu > Au > Ag.

Further insight into the adsorption mechanism can be obtained by decomposing the total interaction energy to the interaction energies of the different parts of S^up^ with the metal surfaces (**Table 3**). The results indicated that glycan groups of S^up^ showed the highest interaction energy. This is absolutely in agreement with the analysis of the contacts between the protein and the surface that indicated that the largest contribution to the protein-surface contacts came from the glycans (**Figure 4c**).

**Table 3.** Decomposition of the interaction energy between S^up^ and the metal surfaces to the different parts of the protein.

|  | Protein | RBD | Glycans | Total |  |
| --- | --- | --- | --- | --- | --- |
| Interaction<br>energy/kJ.mol<sup>-1</sup> | -416.73±20.83 | -219.56±11.65 | -1677.22±85.41 | -2313.51±117.89 | Ag |
|  | -611.36±17.07 | -409.71±24.68 | -2189.91±106.44 | -3210.98±148.19 | Au |
|  | -3523.86±134.07 | -2458.25±129.55 | -7582.49±353.51 | -13564.60±617.13 | Cu |

##### Conformational changes

To understand the conformational changes of S^up^, we have calculated its RMSD relative to the crystal structure. Hydrogen atoms and glycans were excluded from this calculation since they are extremely labile and their fluctuations do not reflect changes in the protein conformation. As shown in **Figure 6a**, the RMSD of the simulated systems initially increases with time, reaching equilibrium values after about 75, 25 and, 150 ns for Ag, Au, and Cu, respectively. As shown in **Figure 6a**, the RMSD values of S^up^ averaged over equilibrium configurations were about 1.48±0.46, 2.00±0.51, and 1.14±0.30 nm for the adsorption onto the surfaces of Ag, Au and Cu, respectively. Hence, it can be concluded that the structural changes of S^up^ adsorbed onto the metal surfaces follow Au > Ag > Cu. Similar behavior has been seen for the adsorption of peptides to Cu and Au surfaces before [104].

**Figure 6.**
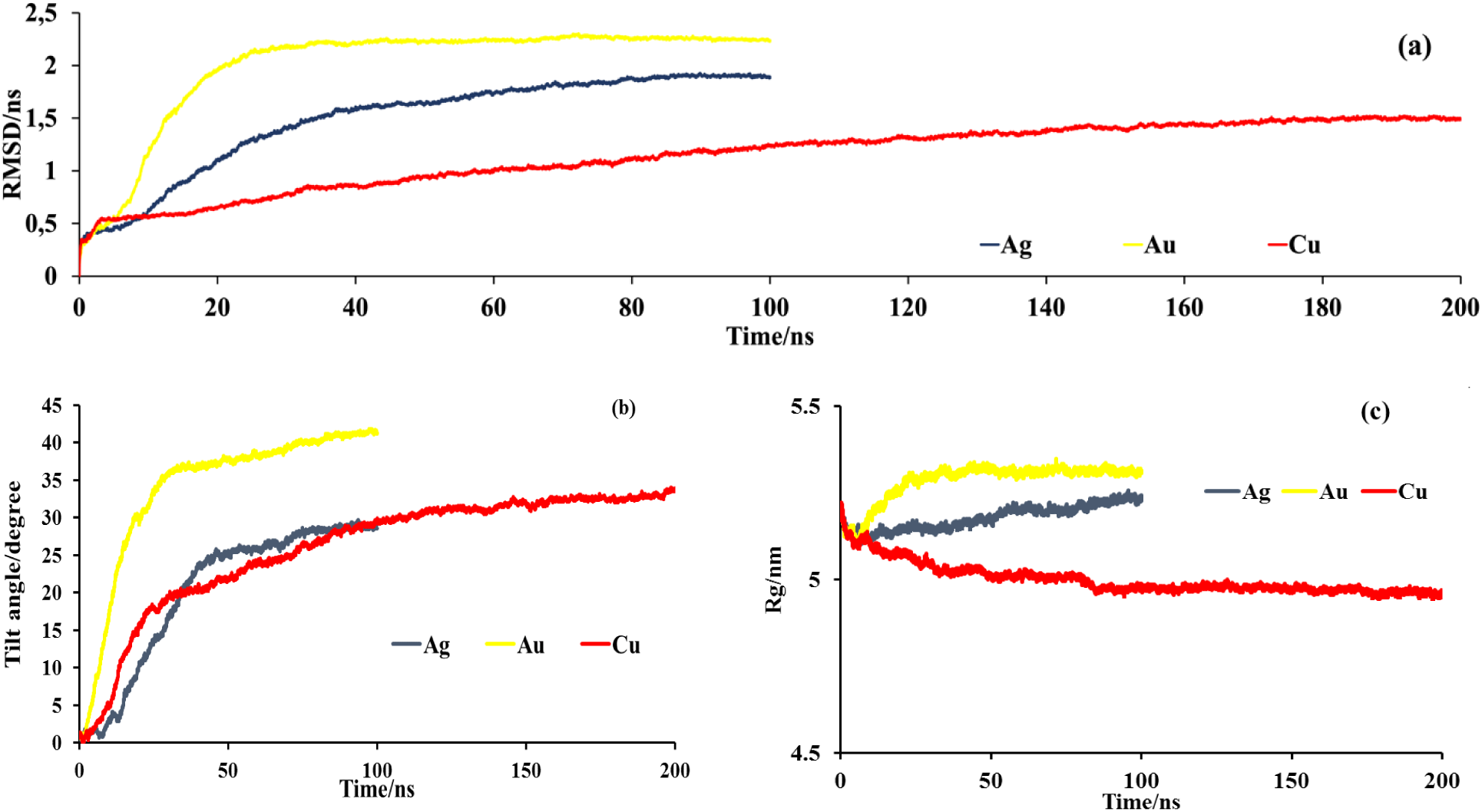
(a) RMSD of all backbone carbon atoms of S^pro^, (b) Tilt angle between S^pro^ and the z axis (the axis perpendicular to the surface) as a function of time, and (c) Time evolution of R_g_ of S^pro^ during its interaction with metal surfaces.

**Figure 6b** represents the tilt angle of the protein. The tilt angle between the major axis of S^up^ and the z-axis (the axis perpendicular to the surface) was computed using the “gmx bundle” module of the gromacs. This module analyzes bundles of axes and reads two index groups and the centers of mass of these groups define the tops and bottoms of the axes. As it is clear from this figure (and also **Figures 3 and S3**), the final equilibrium angles of S^up^ are about 28.53°, 41.24°, and 33.88° onto the Ag, Au, and Cu surfaces, respectively. It means that the direction of S^up^ shows more changes during interaction with the surface of Au. This observation is consistent with the above-mentioned results that show the larger RMSD changes of the protein during the interaction with Au. Therefore, it can be concluded that among the studied metals, Au has the highest ability to change the protein structure and Cu has the highest affinity for S^up^. Both of these characteristics are crucial to designing a new generation of virucidal materials/coatings. In fact, among the three studied metals, Cu is more likely to accumulate virus particles, but Au is more likely to impact on the virus structure.

Radius of gyration (R_g_) can be considered as an index of compactness, stability, and folding state of a protein. As it can be seen in **Figure 6c**, the initial R_g_ value of S^up^ is about 5.17 nm. However, during the MD simulation time its value decreases to 4.95 nm on the Cu surface and increases to 5.23 and 5.31 nm on the Ag and Au surfaces, respectively. Indeed, S^up^ loses its compactness during adsorption onto the surface of Au. On the other hand, the stronger interaction (more negative LJ interaction energy) of S^up^ with the Cu surface causes a more compact structure for it that is in good agreement with the conformational entropy concept stated above. Furthermore, the R_g_ values were stabilized at about 75, 25 and 150 ns during the adsorption onto the surface of Ag, Au, and Cu, respectively. This stabilization that was observed for RMSD, as well, indicates that the MD simulation achieved equilibrium thereafter.

##### Secondary structure

Finally, the secondary structure of S^up^ was analyzed using the DSSP module [105]. The result provides the α-helix, β-sheet along with the total secondary structure contents of the protein. It is easy to notice that the main secondary structures of the protein in the presence of the metals remain stable during whole MD simulation time (**Table 4**). Therefore, during the interaction of the protein with the Ag, Au and Cu surfaces, the tertiary structure of the protein has been changed and adjusted in such a manner to stabilize the metal-S^up^ complexes (especially for Cu) but its secondary structures remain stable.

**Table 4.** Changes of secondary structure of S^up^ due to its adsorption onto the metal surfaces.

| Type of 2<sup>nd</sup> structure | % Structure* | | % α-helix | | % β-sheet | |
| --- | --- | --- | --- | --- | --- | --- |
|  | Initial | Final | Initial | Final | Initial | Final |
| **Ag** | 37.2 | 36.6 | 11.9 | 11.4 | 18.4 | 17.8 |
| **Au** | 37.2 | 36.3 | 11.9 | 11.7 | 18.4 | 18.2 |
| **Cu** | 37.2 | 36.3 | 11.9 | 11.4 | 18.4 | 17.8 |

\*Structure = $\alpha$ helix + $\beta$ sheet + $\beta$ bridge + Turn

#### 3.2.2 S^down^ conformation

Detailed analysis of 3D images of SARS-CoV-2 virions has revealed that the proportions of S^up^ and S^down^ are approximately 1:1 [73]. Most of the published molecular dynamics (MD) simulation studies have only investigated the interaction of S^up^ with the surface of the materials, and only a few studies have considered the interaction of S^down^, as well [74]. Given the relevance of both conformations, we also consider the interaction of S^down^ to the metal surfaces. **Figure S4** shows the beginning of the interaction and final configurations of S^down^ on the surface of metals. In **Figure 7** we compare the results for both conformations of S^pro^ for the three metal surfaces (Ag, Au, Cu). The results show that the interaction of S^down^ onto the metal surfaces has similarities but also important differences as compared with the case of the up-conformation. In all cases, it seems that the glycan groups have a dominant role during interaction of the protein with the metal surfaces. However, the S^up^ shows more changes during interaction with metal surfaces (higher tilt and higher contact). The origin of this difference will be investigated in more detail in the next sections.

**Figure 7.**
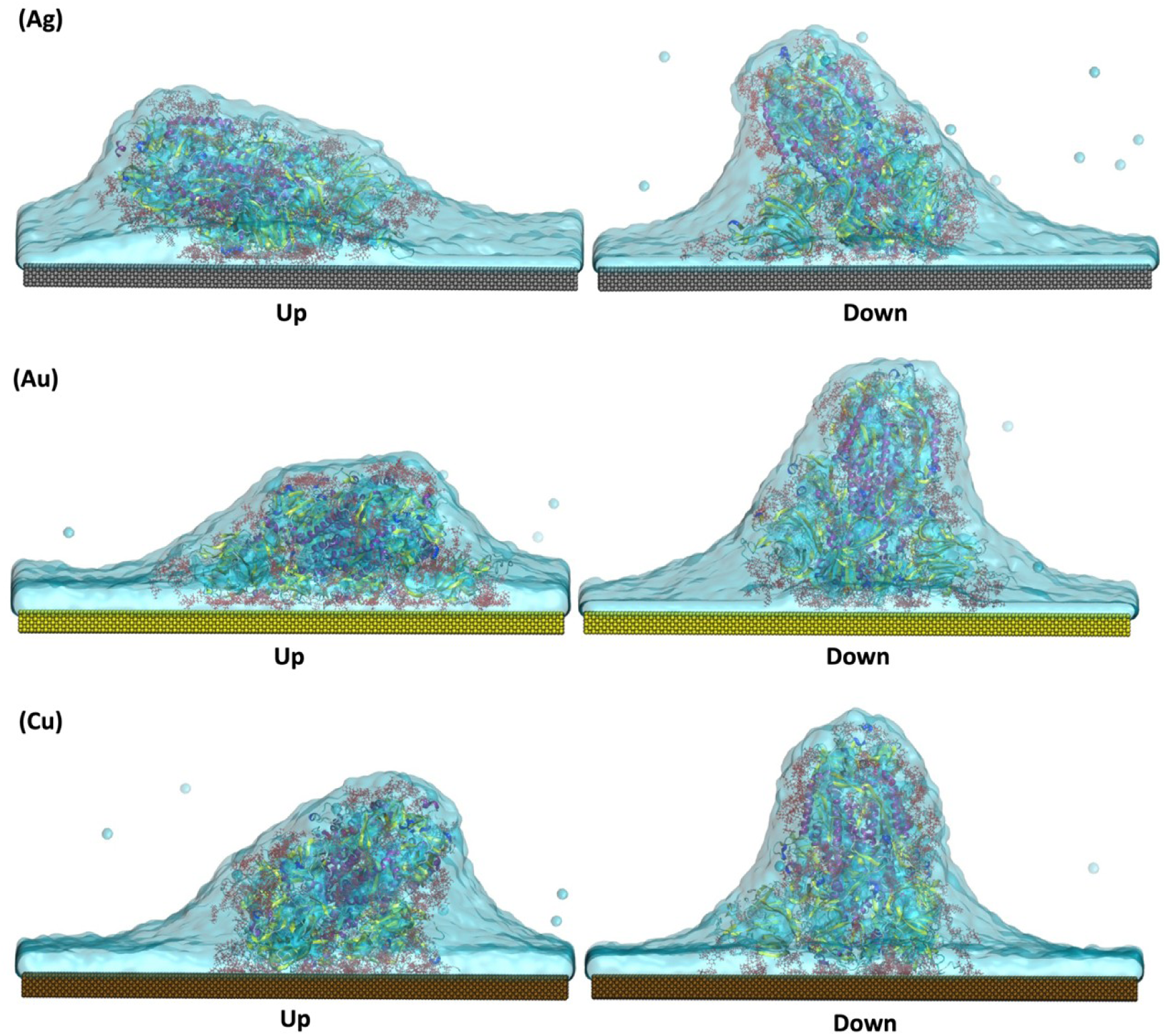
Representative snapshots of the final configurations of the up and down conformations of S^pro^ adsorbed onto Ag, Au, and Cu surfaces. Red CPK representations are glycan groups bound to the spike protein.

**Figure 8** shows that the contact area changes of S^down^ and metal surfaces are almost similar to the changes for S^up^ (**Figure 4**) but with lower values. Similar behavior was also observed for the total number of contacts (data are not shown). Furthermore, Ser, Thr, Asn, Gln and Val amino acid residues as well as glycan groups are responsible for the stability of S^down^ on the surface of metals (similar to S^up^, data are not shown).

**Figure 8.**
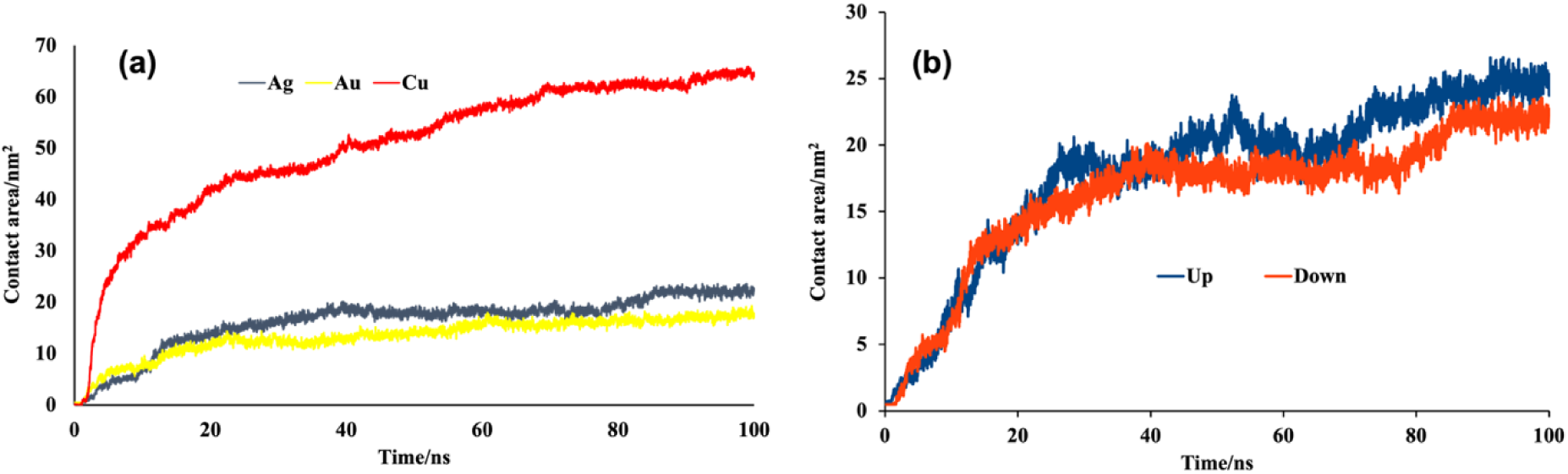
(a) Contact area between S^down^ and metal surfaces as a function of time, (b) comparison between the contact area of S^up^ and S^down^ during the interaction with Ag surface (please see **Figure S6** for Au and Cu surfaces).

RMSF changes were also calculated and shown in **Figure S5**. Although the behavior of this property was almost the same as the observed changes for S^up^ but, as it could be predicted from conformational changes of the proteins on the metal surfaces (**Figure 7**); S^down^ is more stable than S^up^ during the MD simulation time and its amino acid residues shows lower flexibility.

The changes in interaction energy with the evolution of time for S^down^ were also consistent with the contact results discussed above and showed the same behavior as the observed for S^up^. Indeed, these observations also suggest that Cu has the highest affinity for adsorption of the up and down conformations of S^pro^ (**Table 5**) that is, in agreement with the obtained results from contact information. Decomposition of the total interaction energy to the interaction energies of the different parts of S^down^ with the surface of the metal slabs is also shown in **Table 5**. The results indicated that glycan groups of the spike showed the highest interaction energy. This result is in agreement with the observed adsorption mechanisms in **Figure 7**. Furthermore, comparison between the results in **Tables 3 and 5** states that the interaction energy of S^down^ with metal surfaces is lower than the interaction energy for S^up^, as could be expected from higher contact area and total number of contacts of S^up^ (**Figure 8**).

**Table 5.**
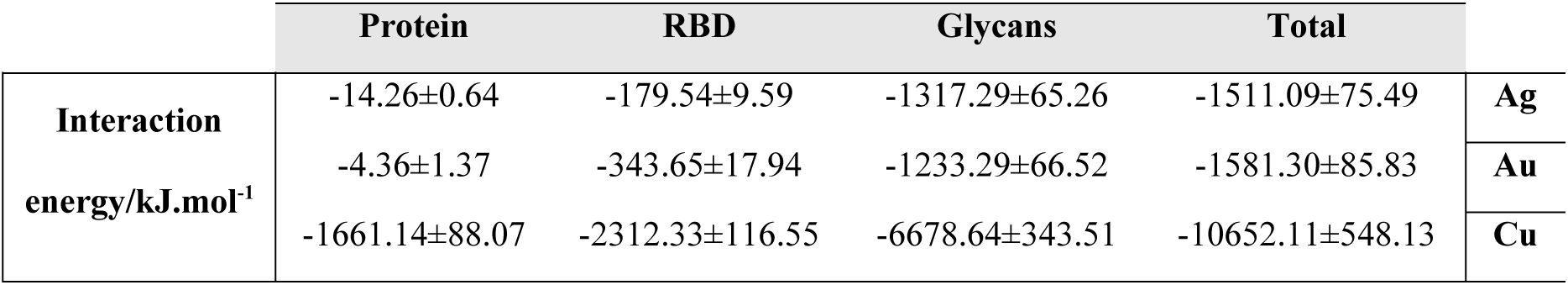
Decomposition of the interaction energy between S^down^ and metal surfaces to the different parts of the protein.

|  | Protein | RBD | Glycans | Total |  |
| --- | --- | --- | --- | --- | --- |
| **Interaction**<br>**energy/kJ.mol<sup>-1</sup>** | -14.26±0.64 | -179.54±9.59 | -1317.29±65.26 | -1511.09±75.49 | **Ag** |
|  | -4.36±1.37 | -343.65±17.94 | -1233.29±66.52 | -1581.30±85.83 | **Au** |
|  | -1661.14±88.07 | -2312.33±116.55 | -6678.64±343.51 | -10652.11±548.13 | **Cu** |

To understand more details about the conformational changes of S^down^, we calculated its RMSD relative to the crystal structure and without hydrogen atoms and glycans, tilt angle and R_g_. The average RMSD values of spike protein were about 0.66±0.14, 0.50±0.07, and 0.50±0.05 nm during the adsorption onto the surface of Ag, Au, and Cu, respectively. Hence, the structural change of S^down^ adsorbed onto the Ag surface is slightly higher than in the two other cases. This observation is consistent with the lowest interaction energy of S^down^ during the adsorption on to the Ag surface. Furthermore, analysis of **Figures 9a and 9b** shows that the RMSD of S^down^ on the Ag surface is less than its value for S^up^. This same behavior is also seen on the surface of Au and Cu (data not shown).

**Figure 9.**
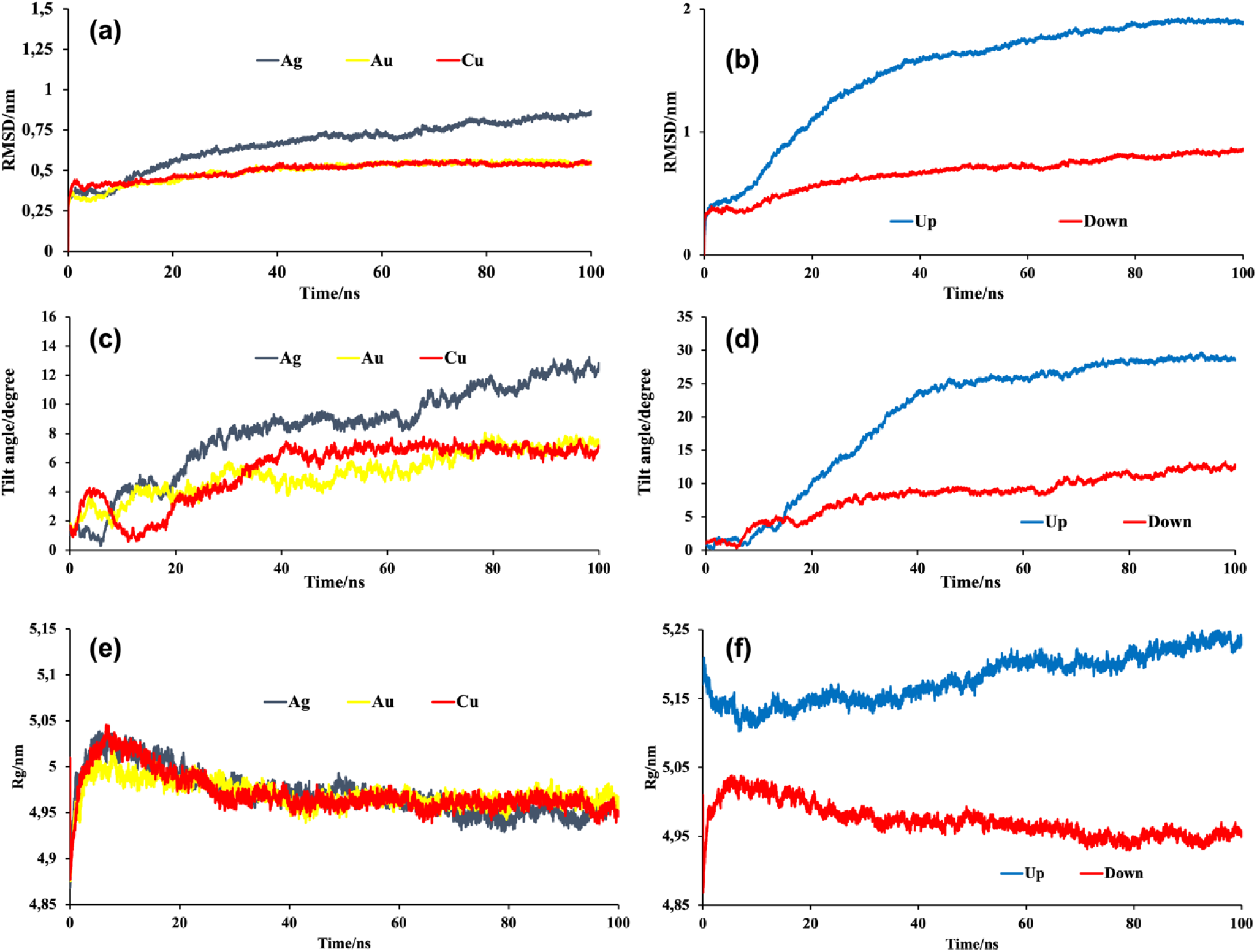
(a) RMSD of all backbone carbon atoms of S^down^. (b) Comparing S^up^ and S^down^ during the adsorption onto the Ag surface based on their RMSD, (c) Tilt angle between S^down^ and the z axis (the axis perpendicular to the surface) as a function of time, (d) Comparing S^up^ and S^down^ based on their tilt angle during the adsorption onto the Ag surface, (e) Time evolution of R_g_ of S^down^ during its interaction with metal surfaces and (f) Comparing the R_g_ values of S^up^ and S^down^ during the adsorption onto the Ag surface. (The results of comparison between S^up^ and S^down^ on the Au and Cu surfaces represented the same behavior as Ag, data not shown.)

**Figure 9c** represents the tilt angle of the protein axis with respect to the z-direction. As it is clear from this figure (and also **Figure 7**), the final equilibrium angles are about 12.77°, 7.47° and 7.04° for the Ag, Au, and Cu, respectively. Although this observation is consistent with the above-mentioned results that showed the larger RMSD changes of the protein during the interaction with Ag, it could be also concluded that in comparison to S^up^, S^down^ does not substantially changes its orientation during the interaction with the surfaces (**Figure 9d**).

As it can be seen in **Figures 9e and 9f**, the initial R_g_ value of S^down^ is about 5.01 nm, which is lower than its value for S^up^. Moreover, during MD simulation time, its R_g_ value decreases to about 4.95 nm on the surface of all three metals. Indeed, S^down^ goes to a more compact structure during adsorption onto the metal surfaces.

Finally, we analyzed the secondary structural changes of S^down^ during the interaction with the metal surfaces, but no considerable changes were observed (similar to S^up^ (**Table 4**), data not shown).

### 3.3. Interaction of S^pro^ with different surfaces: a comparison of MD simulation results

In addition to the metal surfaces (Ag, Au and Cu), we have also investigated a variety of other hard (graphite and cellulose) and soft (different skin models) materials for probing their interaction with spike protein in our research group using the same methodology considered here [69, 70]. Hence, it could be interesting to compare our presence results with the previous ones to make a general scheme about the surface-type effect on adsorption of S^pro^. The results for all the studied systems are collected in **Table 6**.

**Table 6.** Comparison of MD simulations results for interaction of the S^pro^ and different type of surfaces. All the data are corresponding to the up conformation of S^pro^. *”Sebum” means a model of the sebaceous outer layer of human skin, “Stratum corneum” is the nonsebaceous outer layer of human skin, and “POPC” refers to a POPC phospholipid bilayer and it can be understood as a generic model for soft matter.

|  | Cellulose | Graphite | Sebum* | Stratum corneum* | POPC* | Ag | Au | Cu |
| --- | --- | --- | --- | --- | --- | --- | --- | --- |
| **No. contacts** | 51±2 | 96±2 | 87±6 | 180±6 | 158±8 | 24±1 | 14±1 | 94±4 |
| **No. Hbonds** | 18±4 | - | 11±3 | 74±8 | 75±8 | - | - | - |
| **RMSD/Å** | 8.1±0.2 | 18.3±0.3 | 3.8±0.2 | 6.8±0.1 | 5.7±0.2 | 14.8±0.66 | 20.0±0.91 | 11.4±0.54 |
| **Tilt angle/degree** | 45.3±2.3 | 76.4±0.7 | 16.2±0.7 | 83.0±0.5 | 82.9±1.5 | 28.53±0.5 | 41.24±2.1 | 33.88±1.6 |

**Figure 10** represents the RMSD changes of S^pro^ versus the total number of contacts during its interaction with different types of materials we have studied until now. The RMSD and total number of contacts can be considered as indices for the structural changes of the protein in comparison to its crystal structure and affinity of the protein to be adsorbed on the surface of materials, respectively. In this regard, we can imagine four different types of materials based on their interaction with S^pro^. Group (i) corresponds to materials that have a low number of contacts between S^pro^ and the surface and induce a small change in the protein. We characterize them as having RMSD below 10 Å and total number of contacts below 100. These materials have a low affinity for S^pro^ and are not able to change its structure. Hence, they can be considered as materials that have no special effect on infective viral particles (Cellulose and Sebum in our studies). Group (ii) corresponds to materials that induce a substantial change in the protein structure but have a low affinity for binding S^pro^. Quantitatively, we define them as having RMSD above 10 Å and total number of contacts below 100. Hence, they can be considered as materials that may inactivate SARS-CoV-2 virus but are less likely to accumulate infective viral particles (Graphite and metals in our studies). Group (iii) is assigned to RMSD below 10 Å and number of contacts above 100: these materials have a high affinity for S^pro^ but are not able to change its structure. Therefore, this group consists of materials with the ability to capture and accumulate the infective viral particles, but they cannot inactivate them (POPC and Stratum corneum in our studies). This group might be able to inhibit virus transmission. Finally, group (iv) corresponds to RMSD and number of contacts above 10 Å and 100, respectively: these materials not only have a high affinity for S^pro^ but also that have a high ability to change the protein structure, as well. Hence, materials classified in this group may be the most suitable for use as virucidal materials or main components of personal protective equipment [106, 107]. Unfortunately, we have not identified any material belonging to this class until now.

**Figure 10.**
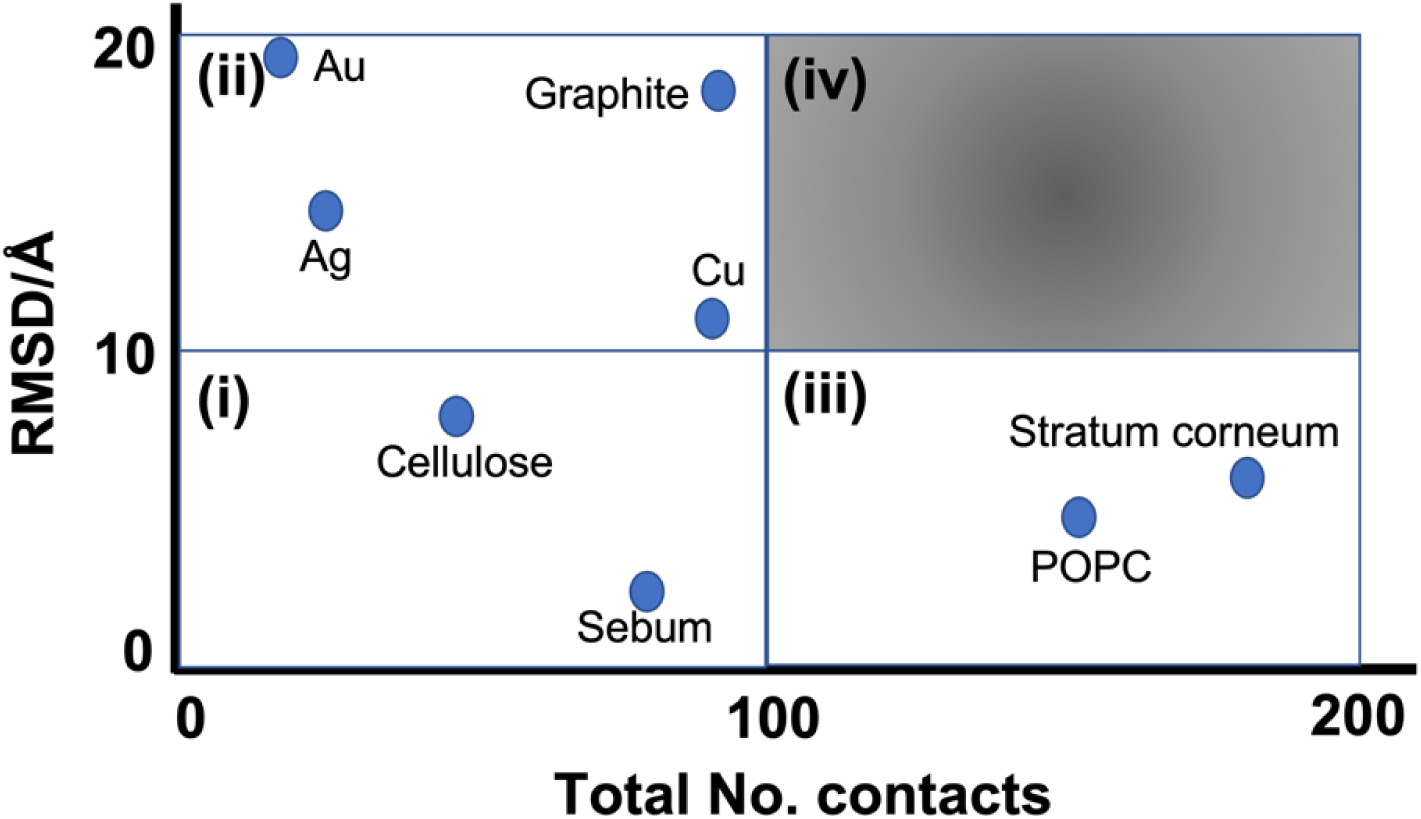
The RMSD changes of S^pro^ versus total number of contacts during its interaction with different types of materials we have studied till now. The i-iv classifications are described in the main text.

The scheme in **Figure 10** can be used to classify the available and future results for interaction of S^pro^ with different materials in a rational manner.

## 4. CONCLUSIONS

In the present work we investigated the interaction of the S1 subunit of S^pro^ (as responsible part of the virus to interact with environment) with coinage metals (Ag, Au, and Cu) using the all-atom MD simulation method. Both of the protein conformations (S^up^ and S^down^) were considered in this study due to their equivalent proportion in SARS-CoV-2 virions. First, we verified the used water model and force field parameters for the metals by simulation of their wetting behavior. The analysis of the trajectories for spike S^pro^-metal complexes revealed that both conformations undergo similar mechanisms to be adsorbed on the surface of metals. The hydrophobic and vdW interactions of metals with the hydrophobic residues and glycan groups of S^pro^ were the main driving forces for adsorption of the protein. Interestingly, the spike protein in the S^up^ conformation has more contacts with the metal surfaces than in the S^down^ conformation. Following the changes in interaction energies showed that although the adsorption mechanisms of S^up^ and S^down^ are similar, their interaction energies during the adsorption onto the metals are different and the affinity of metals for S^up^ is higher than S^down^. On the other hand, Cu shows the highest (most negative) interaction energy among the studied metals. It seems that vdW parameters (ε, σ and their ratio) of metals are important factors to determine the affinity of the metal surfaces for S^pro^ and Cu with the highest ε/σ ratio, and the lowest ε value (among the studied metals) has the highest interaction energy with S^pro^. Investigation of RMSD, tilt angel and R_g_ revealed that the ability of metals to disrupt the 3D conformation of S^up^ follows as Au > Ag > Cu and for S^down^ this behavior changes as Ag > Au > Cu. The lowest conformational changes on the surface of Cu could be related to the strongest interaction of S^pro^ with the Cu surface that hindered the protein motion. Indeed, it could be stated that conformational entropy may play an important role during the adsorption process. Although the tertiary structural changes for S^up^ were more conspicuous than S^down^, adhesion to the metals did not cause any tangible secondary structural changes of the protein. Evaluation of the protein residues mobility (RMSF) showed that S^down^ is more stable than S^up^ during the MD simulation time and its amino acid residues show lower flexibility than the amino acid residues of S^up^. Finally, we complemented our present and previously published results to classify the studied hard and soft materials based on their affinity for S^pro^ and their ability to change its conformation. Based on our classification, polymeric materials like cellulose have no special effect on infective viral particles but carbon-based materials like graphite and the present investigated metals may inactivate SARS-CoV-2 virus. Also, POPC and Stratum corneum have the ability to capture and accumulate the infective viral particles but cannot inactivate them. Moreover, among the three studied coinage metals, Cu is more likely to accumulate virus particles, but Au tends to destroy the virus structure.

Our results can shed the light to investigate the fundamental physico-chemical aspects of the virus-surface interaction in order to identify which factors may make a surface prone to virus adhesion or make it virucidal. Furthermore, the presented classification provides a general view that might pave the way for developing a new generation of virucidal materials/coatings based on coinage metals.

## DATA AND SOFTWARE AVAILABILITY

Fully glycosylated structures of the S1 subunit of SARS-CoV-2 spike protein were taken from CHARMM-GUI archive (https://www.charmm-gui.org/?doc=archive&lib=covid19). Specific open access software from third parties were also used: GROMACS version 2019.3 (https://www.manual.gromacs.org/documentation/2019.3/download.html/), VMD1.9 (http://www.ks.uiuc.edu/Research/vmd/) and Avogadro (https://www.avogadro.cc). Graphical abstract was created with http://www.BioRender.com/. Input and output files are available at https://github.com/soft-matter-theory-at-icmab-csic (and also from the corresponding author upon request).

## ASSOCIATED CONTENT

### Supporting Information

Fully glycosylated structures of the S1 subunit of SARS-CoV-2 spike protein, RMSD evolution of water droplet, Final snapshots of S^pro^ adsorbed onto the metals, Representative snapshots of the down conformation of spike protein adsorbed onto the metals, Comparison between the RMSF of up and down conformations of S^pro^, Comparison between the contact area of up and down conformations of S^pro^.

## AUTHORS INFORMATION

**Mehdi Sahihi**-Institut de Ciencia de Materials de Barcelona (ICMAB-CSIC), Campus de la UAB, E-08193 Bellaterra, Barcelona, Spain, https://orcid.org/0000-0003-2923-1833

**Jordi Faraudo**-Institut de Ciencia de Materials de Barcelona (ICMAB-CSIC), Campus de la UAB, E-08193 Bellaterra, Barcelona, Spain, https://orcid.org/0000-0002-6315-4993

## Supporting information

Further details and additional figures for our simulations

## ACKNOWLEDGMENT

This work was supported by Grant PID2021-124297NB-C33 funded by MCIN/AEI/ 10.13039/501100011033 and, as appropriate, by “ERDF A way of making Europe”, by the “European Union” or by the “European Union NextGenerationEU/PRTR” and by the “Severo Ochoa” Program for Centers of Excellence in R&D (CEX2019-000917-S) awarded to ICMAB. We thank the Spanish national supercomputing network (BSC-RES) for the award of computer time at the Minotauro supercomputer. M Sahihi is supported by the European Union Horizon 2020 research and innovation programme under Marie Sklodowska-Curie Action Individual Fellowship Grant Agreement No. 101026158.

