## Supplementary material for "Computer Simulation of the interaction between SARS-CoV-2 Spike Protein and the Surface of Coinage Metals": Further details and additional figures for our simulations

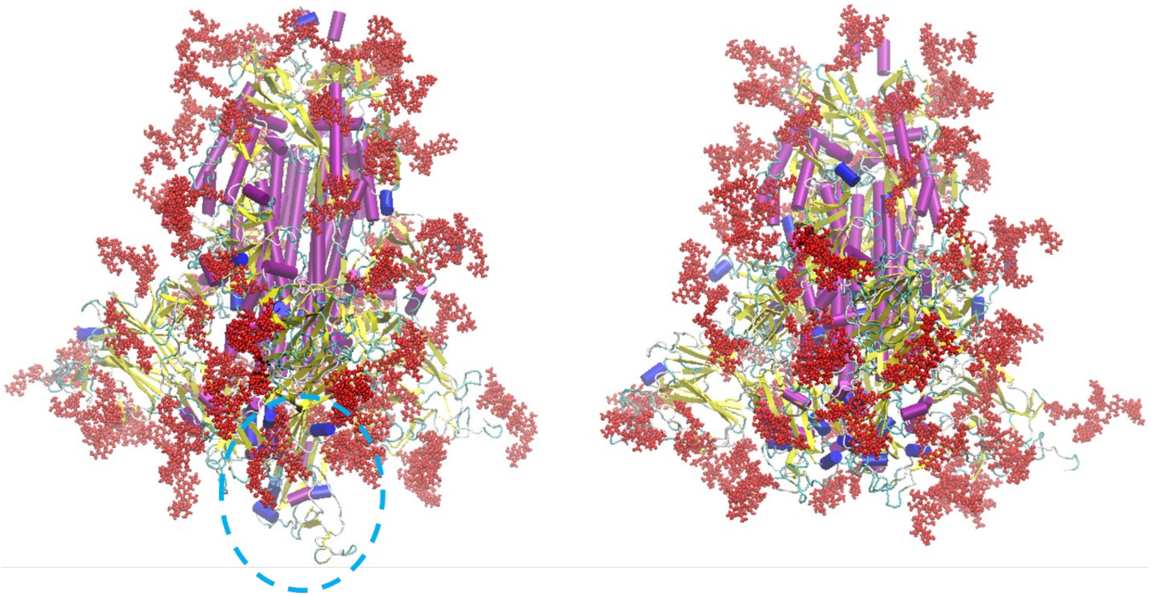


**Figure S1.** Fully glycosylated structures of the S1 subunit of $S^{pro}$ were taken from CHARMM-GUI archive (PDB IDs: 6VSB and 6VXX for up and down conformations, respectively. Glycan groups are shown by red CPK representation.

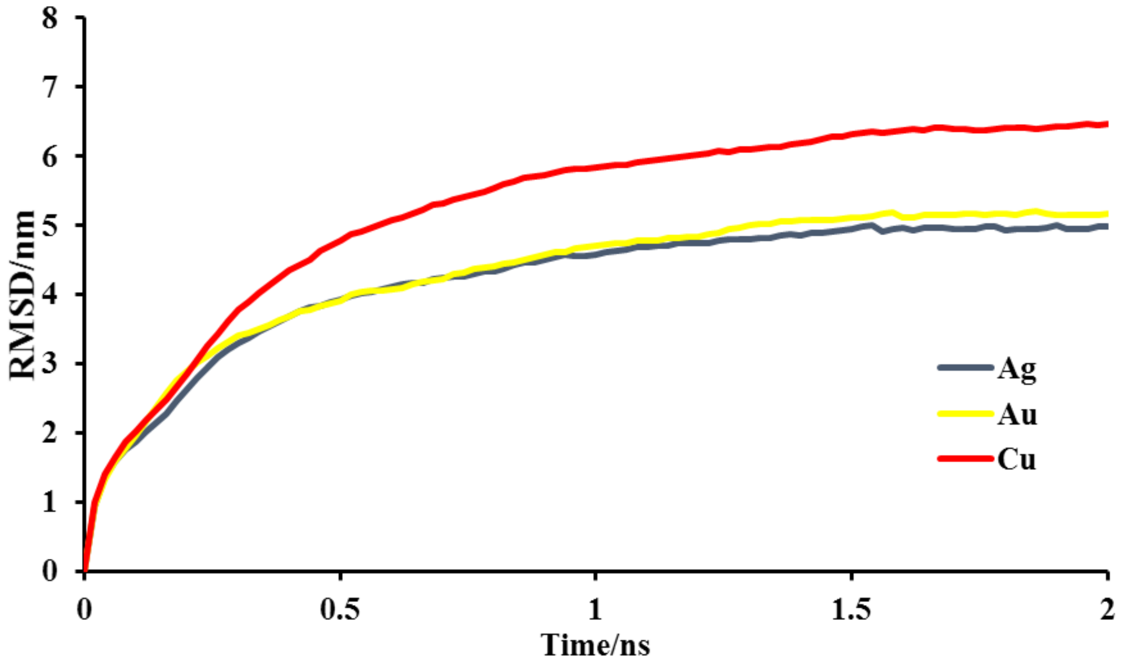


**Figure S2.** RMSD evolution of water droplet during the interaction with metal surfaces.

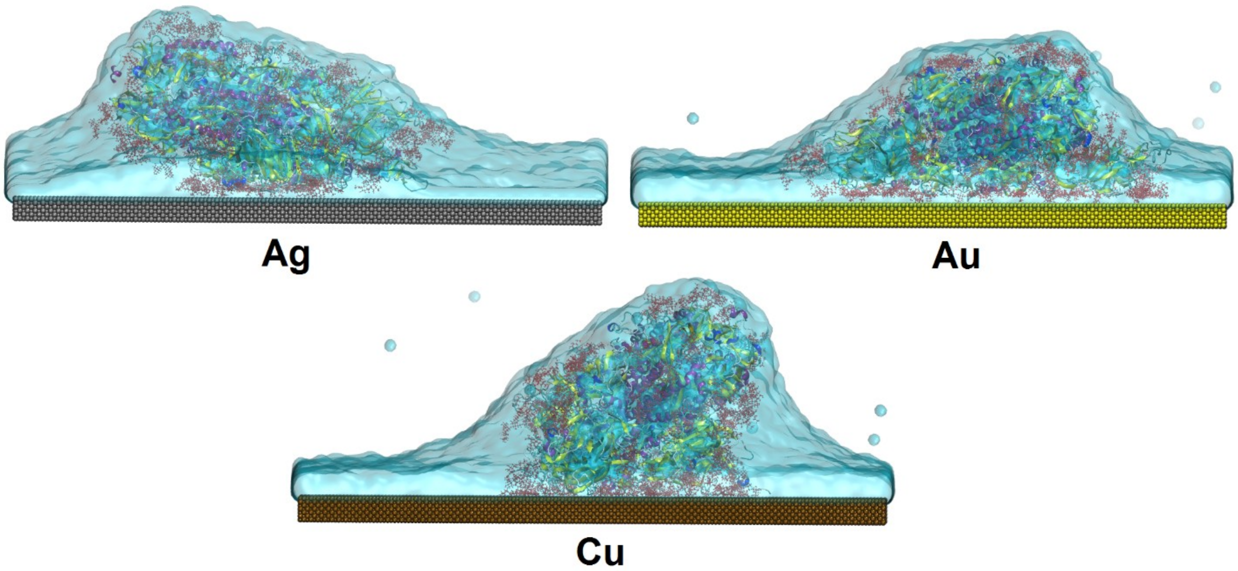


**Figure S3.** The final snapshots of $S^{pro}$ adsorbed onto Ag, Au, and Cu surfaces.

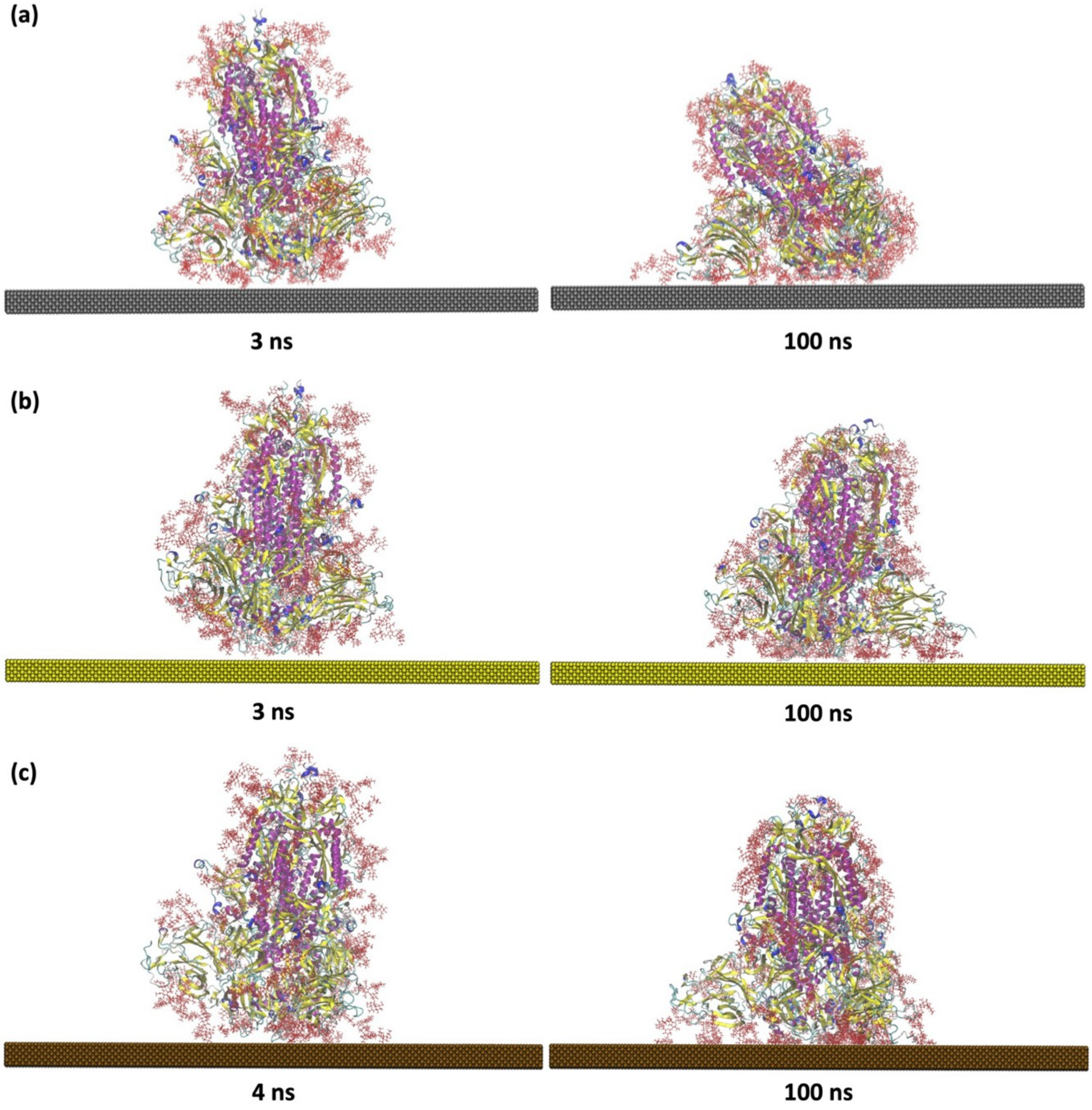


**Figure S4.** Representative snapshots of the down conformation of spike protein adsorbed onto (a) Ag, (b) Au, and (c) Cu surfaces. The solvation water is not shown in the figure to simplify the visualization (**Figure 7** includes solvation water). The number and type of spike amino acid residues (in the final time frame of the trajectories) in contact with metals are shown in **Figure 8**.

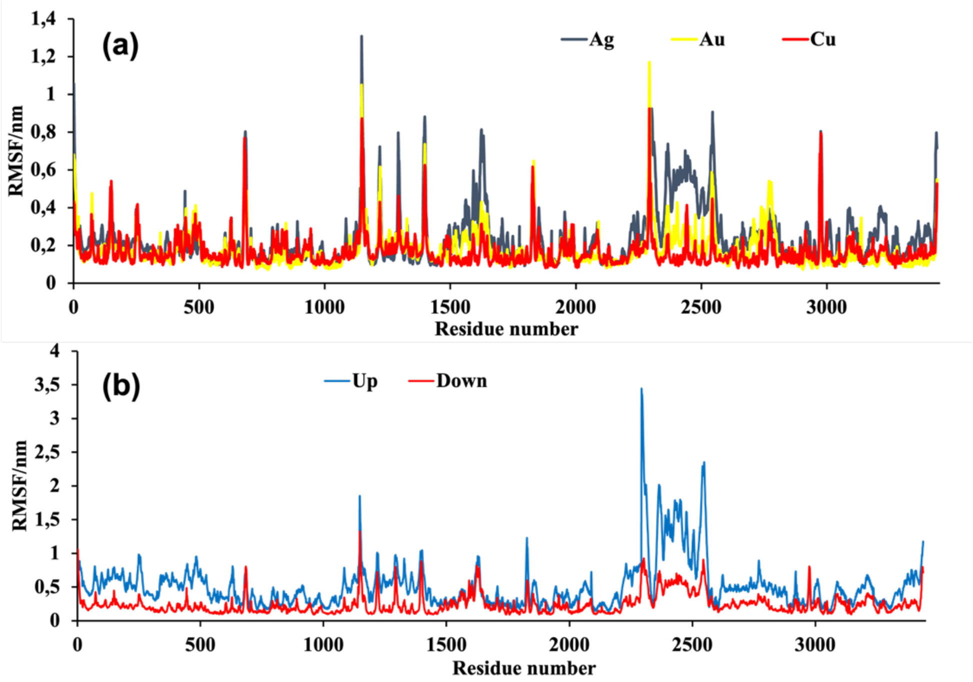


**Figure S5.** (a) RMSF of $S^{down}$ amino acid residues on the surface of metals, (b) Comparing the $S^{up}$ and $S^{down}$ conformations based on their amino acid residues flexibility during the interaction with the Ag surface. (The results for Au and Cu surfaces represent similar behavior and are not shown).

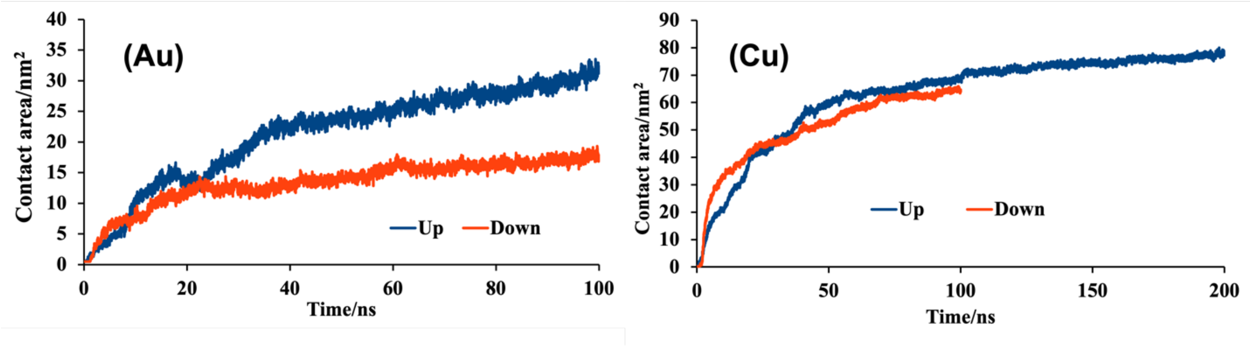


**Figure S6.** Comparison between the contact area of up and down conformations of spike protein during the interaction with Au and Cu surfaces.
